# The carboxyl-terminal processing proteases Prc and CtpA modulate cell-surface signalling activity and *Pseudomonas aeruginosa* virulence

**DOI:** 10.1101/2023.03.13.532344

**Authors:** Joaquín R. Otero-Asman, Ana Sánchez-Jiménez, Karlijn C. Bastiaansen, Sarah Wettstadt, Cristina Civantos, Alicia García-Puente, Wilbert Bitter, María A. Llamas

## Abstract

Cell-surface signalling (CSS) is a signal transfer system of Gram-negative bacteria used to detect extracellular signals and modulate gene transcription in response. These three-protein systems are formed by an outer membrane receptor, a cytoplasmic membrane-embedded anti-σ factor and a cytosolic extracytoplasmic function σ factor (σ^ECF^). In absence of an inducing signal, the anti-σ factor binds to and keeps the σ^ECF^ factor sequestered, thus preventing its interaction with the RNA polymerase and the transcription of signal response genes. Presence of the signal triggers a signalling cascade that extends from the outer membrane to the cytosol and results in σ^ECF^ factor activation. Recently, we and others have reported that CSS σ^ECF^ factor activation requires the regulated and sequential proteolysis of the cognate anti-σ factor, and the function of the Prc and RseP proteases. However, many features of this proteolytic cascade are still unclear. In this work, we have identified another protease that modulates CSS activity, namely the periplasmic carboxyl-terminal processing protease CtpA. We show that both CtpA and the previously identified protease Prc control CSS activation by modulating the levels of the anti-σ factor. CtpA functions upstream of Prc in the proteolytic cascade and seems to prevent the Prc-mediated proteolysis of the CSS anti-σ factor. Importantly, using zebrafish embryos and the A549 cell line as hosts, we show that mutants in the *rseP* and *ctpA* proteases of the human pathogen *Pseudomonas aeruginosa* are considerably attenuated in virulence while the *prc* mutation increases virulence likely by enhancing the production of outer membrane vesicles. Because proteases are druggable proteins, the identification of regulatory proteases involved in *P. aeruginosa* virulence holds promise for the development of novel strategies to fight this clinically relevant pathogen.

## Introduction

In the human pathogen *Pseudomonas aeruginosa*, extracytoplasmic function sigma (σ^ECF^) factors control important biological functions required for bacterial survival and colonization of the host [1]. Activity of several *P. aeruginosa* σ^ECF^ factors is controlled through a signal transduction cascade known as cell-surface signalling (CSS) that together with an σ^ECF^ factor involves an outer membrane receptor and a membrane-embedded anti-σ factor [2] (Fig. 1). The CSS receptor belongs to the TonB-dependent transporter (TBDT) family and is usually involved in both signal transduction and transport of the inducing signal. Transport occurs through the large ß-barrel C-terminal domain of the protein and needs the energy provided by the TonB-ExbBD system [3] (Fig. 1). Signalling occurs via a small N-terminal domain located in the periplasm of the bacteria that interacts with the CSS anti-σ factor [4] (Fig. 1). CSS anti-σ factors typically are single-pass cytoplasmic membrane proteins that contain a large periplasmic C-domain and a short cytosolic N-domain known as <u>a</u>nti-<u>s</u>igma <u>d</u>omain (ASD). While the C-domain receives the signal from the receptor, the N-domain binds the σ^ECF^ factor and keeps it sequestered in absence of the inducing stimulus [2]. The signal-responsive protein of the CSS pathway is the σ^ECF^ factor, which upon activation binds to the RNA polymerase (RNAP) and directs it to the promoter of the signal response genes initiating gene transcription.

**Fig. 1.**
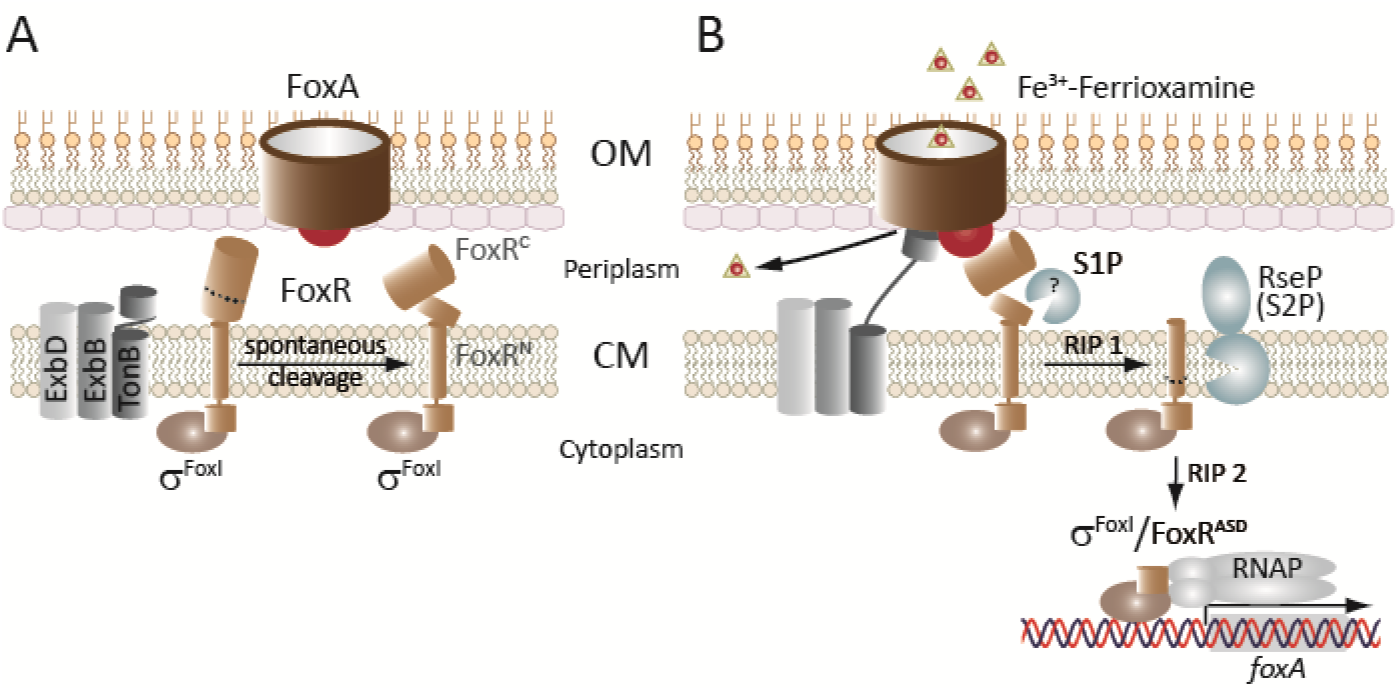
Model of the proteolytic cascade controlling CSS activation based on the *P. aeruginosa* Fox system. **(A)** The three components of the CSS system—receptor (FoxA), anti-σ factor (FoxR), and σ^ECF^ (σ^FoxI^)—and the TonB-ExbBD complex are shown. Prior to signal recognition, the *P. aeruginosa* FoxR anti-σ factor undergoes a spontaneous cleavage that produces two functional N- and C-domains that interact with each other in the periplasm and are both required for proper anti-σ factor function. **(B)** Recognition of the siderophore ferrioxamine by the FoxA receptor produces the interaction of the TonB protein with FoxA enabling the energy coupled uptake of ferrioxamine. Binding of ferrioxamine to FoxA also promotes the interaction of the signalling domain of FoxA (FoxA^SD^, red ball) with FoxR^C^. This event triggers the regulated intramembrane proteolysis (RIP) of the FoxR^N^ domain by the action of (at least) two proteases: a (still unidentified) site-1 protease (S1P) and the site-2 RseP protease (S2P). This results in the release of σ^FoxI^ into the cytoplasm bound to the anti-sigma domain of FoxR (FoxR^ASD^). In several CSS pathways, including the *P. aeruginosa* Fox pathway, the ASD persists and is required for σ^ECF^ activity, having thus pro-σ activity. Although not experimentally demonstrated yet, this domain likely forms part of the transcription complex. Among other genes, σ^FoxI^ promotes the transcription of the *foxA* receptor gene. OM, outer membrane; CM, cytoplasmic membrane; RNAP, RNA polymerase. Adapted from [2] with the findings from [9–11, 17].

CSS systems are extensively present in bacteria of the *Pseudomonas* genus, especially in *P. aeruginosa, P. putida* and *P. protegens*, in contrast to for example *Escherichia coli* that only contains one [1, 2]. Signalling systems increase the bacterial ability to detect and survive in many different environments, a characteristic of *Pseudomonas* bacteria [5, 6]. In this genus, CSS is mainly involved in the regulation of iron acquisition by sensing and responding to iron-chelating compounds like siderophores, iron-citrate or heme, but also in the regulation of bacterial competition and virulence [1, 2, 7]. Analysis of *Pseudomonas* CSS systems has revealed that CSS activation in response to the inducing signal requires the targeted proteolysis of the CSS anti-σ factor (Fig. 1B). This occurs through regulated intramembrane proteolysis (RIP), a conserved mechanism in which a transmembrane protein is subjected to several proteolytic steps in order to liberate and activate a cytosolic effector [8]. The RIP of CSS anti-σ factors always involves the site-2 zinc metalloprotease RseP, which cuts within the transmembrane domain of the protein liberating the CSS σ^ECF^ factor into the cytosol (Fig. 1B) [9–12]. RseP substrate recognition and cleavage occurs through size-filtering rather than by the recognition of a specific sequence/motif [13]. In order to create a suitable substrate for RseP, the anti-σ factor needs to be cleaved by a site-1 protease on the periplasmic side. Evidence that a site-1 cleavage of CSS anti-σ factors occurs is the accumulation of a slightly larger fragment than the RseP product in *rseP* mutants [7, 10, 11]. However, the mechanism triggering this cleavage has not been identified yet. Our earlier analyses identified the carboxyl-terminal processing (CTP) serine protease Prc as the protease required for the site-1 cleavage of the IutY protein of *P. putida* [10]. IutY is a unique protein because it contains the σ^ECF^ and anti-σ factor domains fused in a single protein and separated by a transmembrane domain. Prc is also required for activation of archetypal CSS systems [10], but whether or not Prc directly cleaves these anti-σ factors has not been determined yet. This study aimed to shed light on these unknown features of CSS activation and resulted in the identification of a new protease, CtpA, involved in the activation of this signalling cascade.

## Results

### The CTP serine protease CtpA is involved in CSS activation

Our previous results showed that absent of the Prc protease diminished but did not abolished *P. aeruginosa* CSS activity [10]. We hypothesized that this residual activity could be due to the function of another protease in the absence of Prc. *Pseudomonas* species produces a second carboxyl-terminal processing protease known as CtpA, a soluble periplasmic serine protease that in *P. aeruginosa* has been involved in the activation of the σ^SbrI^ factor [14, 15].

To test the effect of this protease in CSS activity, we introduced a *ctpA* deletion in *P. aeruginosa* (ΔctpA mutant) and assayed activity of the Fox and Fiu CSS systems, which respond to the siderophores ferrioxamine and ferrichrome, respectively. Activity of these systems was assayed using the CSS-dependent transcriptional fusions *foxA::lacZ* and *fiuA::lacZ*, containing the promoter region of the *foxA* or *fiuA* CSS receptor gene fused to a promoterless *lacZ* gene. Expression of these constructs completely depends on the activation of their respective CSS pathways [10, 16]. As expected from previous results, lack of Prc reduced the activity of both CSS systems (Fig. 2A). However, lack of CtpA did not reduce but considerably increased the response of the *P. aeruginosa* Fox and Fiu CSS systems to the presence of their cognate inducing siderophore (Fig. 2A). Absence of this protease in *Pseudomonas putida* had a similar effect (Fig. S1), showing a general role for CtpA in CSS activity. In accordance with increased *foxA* expression in the ΔctpA mutant and decreased in the Δprc mutant, production of the FoxA CSS receptor protein in response to ferrioxamine was higher in ΔctpA and lower in Δprc than in the wild-type PAO1 strain (Fig. 2B). Activity of the Fox system in ΔctpA was not affected in the absence of the siderophore (Fig. 2C), indicating that absence of CtpA does not lead to constitutive activation of the CSS pathways. Production of CtpA from a low-copy number plasmid did not affect the activity of the Fox system in the PAO1 wild-type strain but it was able to complement the *P. aeruginosa* ΔctpA mutation (Fig. 2C). This further indicates that CtpA modulates CSS activity. However, CtpA does not seem to compensate for the lack of Prc as initially hypothesized because the *ctpA* mutation resulted in increased instead of decreased CSS activity. Introduction of the *ctpA* deletion into the Δprc mutant did not alter the phenotype of the single Δprc mutant (Fig. 2A and 2B). The lack of the ΔctpA phenotype in the Δprc ΔctpA double mutant suggests that CtpA works upstream of Prc in modulating CSS activity.

**Fig. 2.**
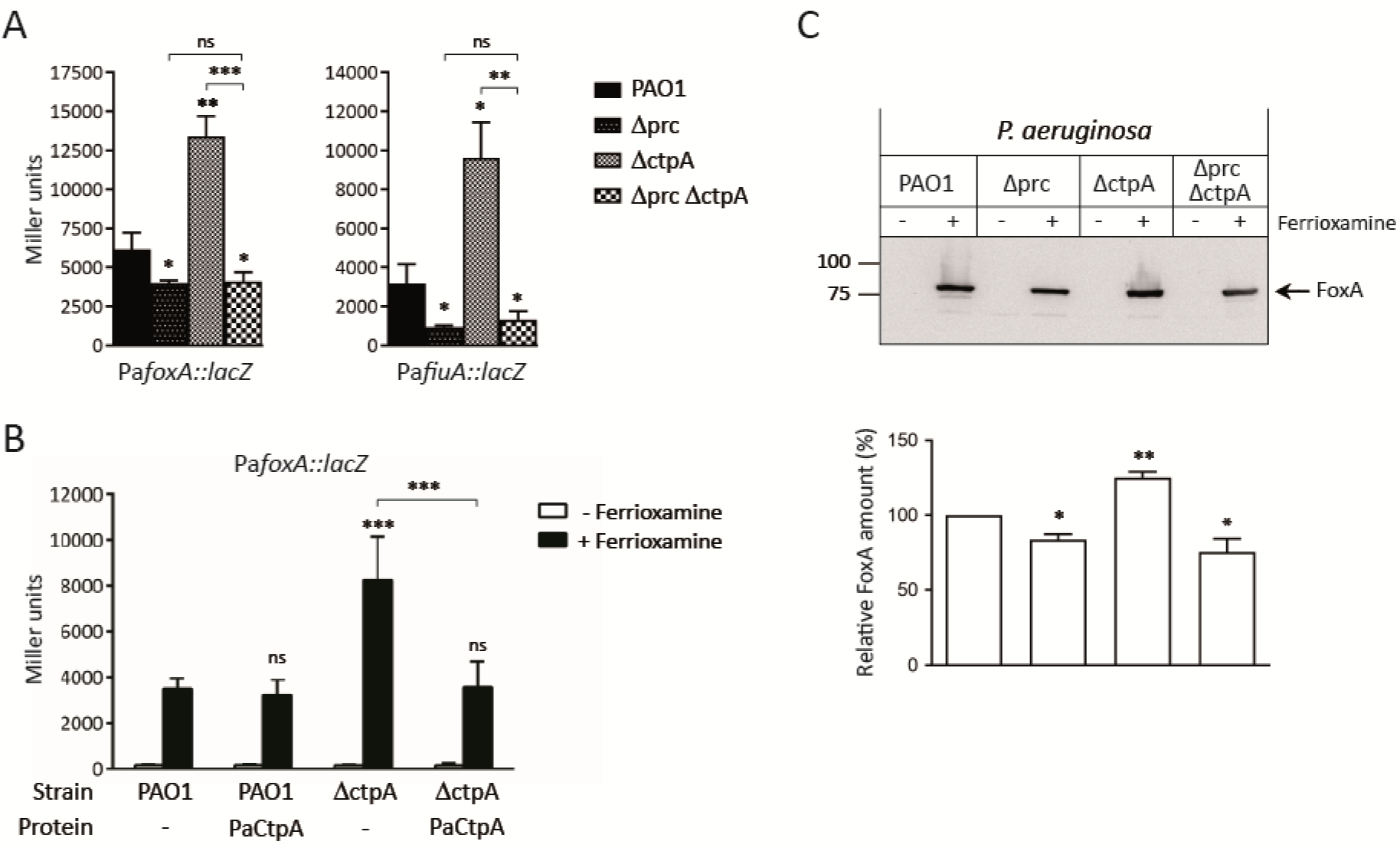
Activity of the *P. aeruginosa Fox* and Fiu CSS systems in *ctpA* mutants. **(A, B)** β-galactosidase activity of the indicated *P. aeruginosa lacZ* fusion gene in the PAO1 wild-type strain and the indicated isogenic mutant. **(A)** Strains were grown under iron-restricted conditions with 1 μM ferrioxamine (Pa*foxA::lacZ*) or 40 μM ferrichrome (Pa*fiuA::lacZ*). In **(B)** strains bear the pBBR1MCS-5 empty (-) or the pBBR/PaCtpA plasmid (Table S1) next to the *PafoxA::lacZ* fusion, and were grown in iron-restricted conditions without (-) or with (+) ferrioxamine. In both panels, data are means ± SD from three biological replicates (N=3). P-values were calculated by two-tailed *t*-test by comparing the value obtained in the mutant with that of the PAO1 wild-type strain in the same growth condition. Comparisons between other strains are indicated by brackets. **(C)** The indicated strains were grown in iron-restricted conditions without (-) or with (+) 1 μM ferrioxamine. Proteins were immunoblotted for FoxA using a polyclonal antibody. Position of FoxA and the molecular size marker (in kDa) is indicated. Blots are representative of three biological replicates (N=3). The graph shows the intensity of the bands obtained with each strain normalized to the PAO1 strain and data are means ± SD from three biological replicates (N=3). P-values were calculated by one-sample *t*-test to a hypothetical value of 100 by comparing the value obtained in the mutant with that of the PAO1 wild-type strain.

### Absence of CtpA increases the amount of the FoxR anti-σ factor protein

We next focused on the identification of the CSS substrate of the *P. aeruginosa* Prc and CtpA proteases. We first considered the CSS anti-σ factor and analysed the *P. aeruginosa* FoxR (PaFoxR) anti-σ factor. To detect PaFoxR, we used N- and C-terminally HA-tagged protein variants. PaFoxR is known to undergo a spontaneous cleavage that leads to a ~22 kDa N-domain (FoxR^N^) and a ~16 kDa C-domain (FoxR^C^) (Fig. 1) [11, 17]. Both domains were detected by Western-blot in considerably higher amounts than the full-length protein (FoxR) (Fig. 3A and 3B, PAO1 strain). In response to ferrioxamine, FoxR^N^ is proteolytically processed by RIP through the action of an unknown site-1 protease and the RseP site-2 protease (Fig. 1B) [11]. RseP cuts within the transmembrane domain of FoxR^N^ and generates FoxR^ASD^ (Fig. 1B), a domain that was detectable upon ferrioxamine induction (Fig. 3A). Lack of CtpA did not affect the RIP of FoxR^N^ and the observed protein bands were obtained in the wild-type and ΔctpA strains (Fig. 3A). However, the amount of the FoxR^N^ domain, and especially that of the FoxR^ASD^, was higher in the ΔctpA mutant (Fig. 3A). Higher amounts of FoxR^ASD^ usually correlates with increased CSS activity likely because this process produces the liberation and activation of the σ^FoxI^ factor [10, 11], a phenotype also observed in the ΔctpA mutant (Fig. 2). Introduction of the Δprc mutation into the ΔctpA mutant reduced the amount of FoxR^ASD^, which was similar to that obtained in the single Δprc mutant (Fig. 3A). This is in agreement with the lower CSS activity observed in the Δprc ΔctpA double mutant (Fig. 2). Interestingly, the *ctpA* mutation had a considerable effect on the FoxR^C^ domain, which was hardly detected in this mutant, especially in presence of ferrioxamine (Fig. 3B). The amount of FoxR^C^ in the Δprc ΔctpA double mutant was similar to that of the wild-type strain (Fig. 3B), confirming that the *prc* mutation abolishes the effect of the *ctpA* mutation. Together, these results suggest that the PaFoxR anti-σ factor is not the substrate of CtpA although the absence of this protease considerably affects the stability of both domains of this protein.

**Fig. 3.**
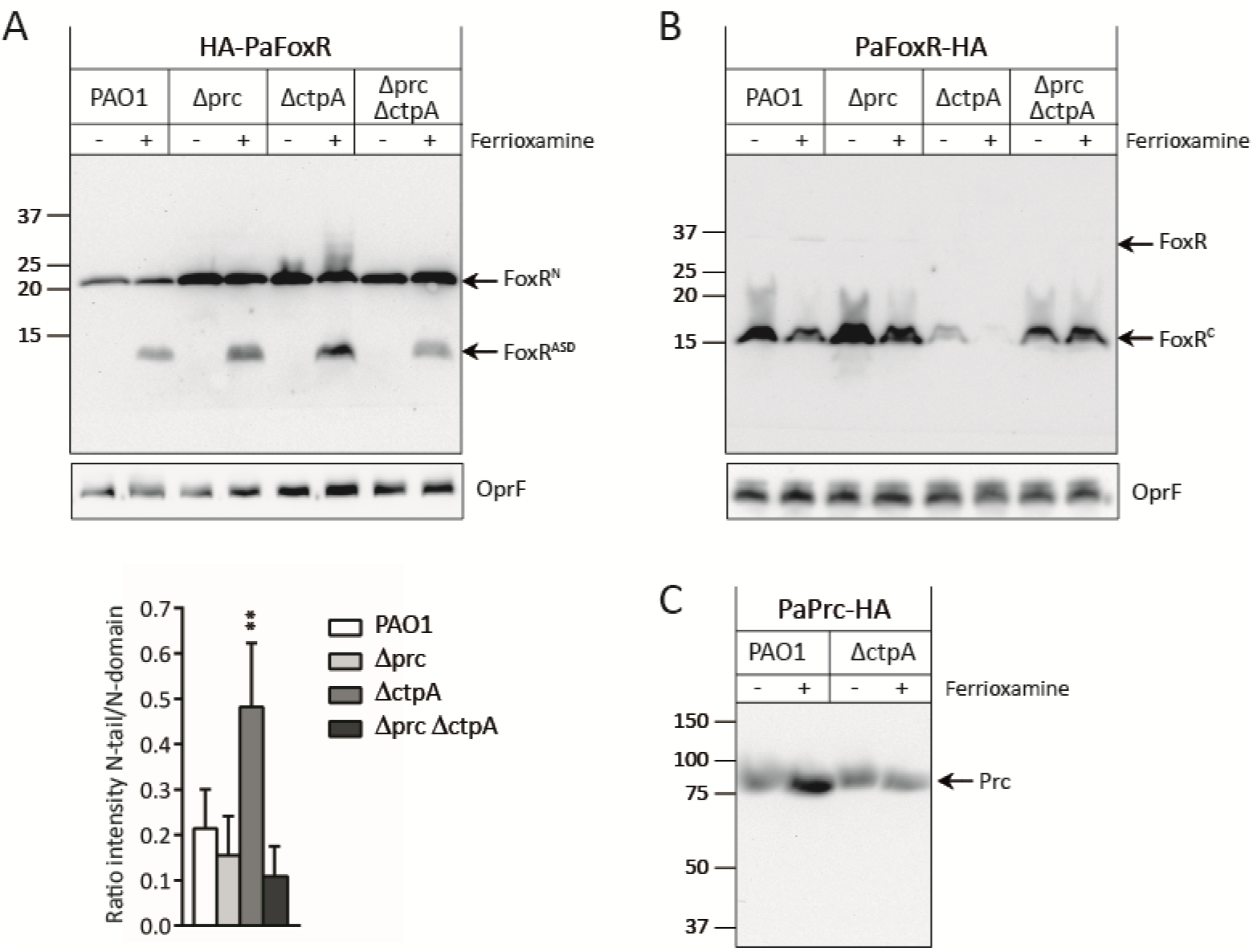
Role of the CtpA protease in processing *P. aeruginosa* CSS components. In all panels, the *P. aeruginosa* PAO1 wild-type strain and the indicated isogenic mutants were grown to late log-phase under iron-restricted conditions and in the absence (-) or presence (+) of 1 μM ferrioxamine. In **(A)** strains express an N-terminally HA-tagged *P. aeruginosa* FoxR protein (HA-PaFoxR), in **(B)** a C-terminally HA-tagged *P. aeruginosa* FoxR protein and in **(C)** a C-terminally HA-tagged Prc protein. Proteins were immunoblotted for HA using a monoclonal antibody. Position of the protein fragments and the molecular size marker (in kDa) is indicated. Presence of the HA-tag adds ~1 kDa to the molar mass of the protein fragments. Blots are representative of at least three biological replicates (N=3). Detection of the OprF protein was used in **(A)** and **(B)** as loading control. The graph in **(A)** shows the ratio between the intensity of the FoxR^ASD^ and the FoxR^N^ domain bands, and data are means ± SD from three biological replicates (N=3). P-values were calculated by two-tailed *t*-test by comparing the value obtained in the mutant with that of the PAO1 wild-type strain.

Because the absence of CtpA lead to an increase of CSS activity in a Prc-dependent manner, we wondered whether Prc itself could be the substrate of CtpA. For this we used an HA-tagged Prc protein. The level of the Prc protease slightly increased in the wild-type strain in presence of ferrioxamine (Fig. 3C, PAO1). This increase was not observed in the ΔctpA mutant (Fig. 3C), which suggests that Prc is more stable when the Fox CSS system is active and that a CtpA-dependent function stabilizes Prc in this condition.

### The proteolytic activities of Prc, CtpA and RseP are required for CSS activation

To further analyse the role of the *P. aeruginosa* Prc and CtpA proteases in the activation of CSS systems, we constructed proteolytically inactive versions of both proteases. For that, we changed the Prc active site protease residue Ser-490 [18] to leucine and the CtpA active site residue Ser-302 [19] to alanine. As control, the active site residue His-21 of the *P. aeruginosa* RseP protease [20] was changed to alanine. Then we assayed the activity of the *P. aeruginosa* Fox CSS system in Δprc, ΔctpA and ΔrseP mutants complemented with either a wild-type Prc, CtpA or RseP protein or with the Prc-S490L, CtpA-S302A or RseP-H21A active site changed versions (Fig. 4A). Activity of the Fox CSS pathway was considerably reduced in the Δprc mutant, highly increased in the ΔctpA mutant, and completely abolished in the ΔrseP mutant (Fig. 4), as shown previously (Fig. 2 and [10]). Complementation of the ΔctpA and ΔrseP mutants with the corresponding wild-type protein restored CSS activity to PAO1 levels, while complementation of the Δprc mutant restored activity only partially (Fig. 4A). The proteolytically inactivated versions of the proteases were unable to restore activity (Fig. 4A). This confirms that the proteolytic activity of the three proteases is required for proper activity of the Fox CSS pathway. Interestingly, the Prc-S490L active site mutant protein was not only unable to complement the Δprc mutation, but also exhibited a dominant negative effect by significantly decreasing the residual *foxA* promoter activity observed in the Δprc background in presence of ferrioxamine (Fig. 4A). Similar results were obtained with active site changed versions of the Prc and RseP proteases of *P. putida* (Fig. S2). While the wild-type versions of the *P. putida* Prc and RseP proteins were able to restore the ferrioxamine- and ferrichrome- induced CSS activity, the proteolytically inactive versions Prc-S485A and RseP-H23A were not (Fig. S2). As observed in *P. aeruginosa*, the proteolytically inactivated version of Prc of *P. putida* significantly decreased the residual *foxA* and *fiuA* promoter activity obtained in the Δprc mutant (Fig. S2), which confirms the dominant negative effect of this protein on CSS activity. Altogether, these results show the importance of the proteolytic action of Prc, CtpA and RseP proteases in activating CSS pathways.

**Fig. 4.**
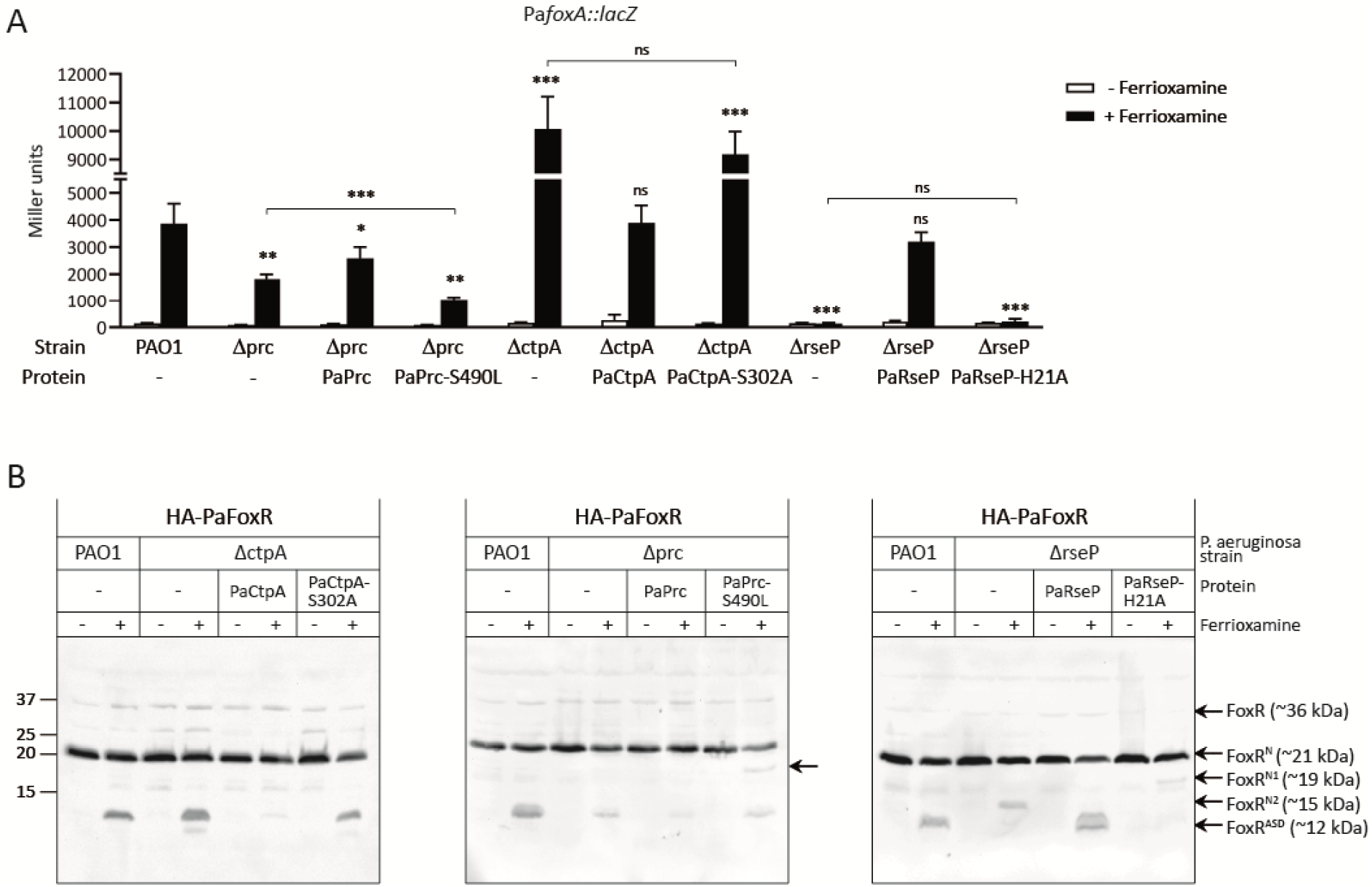
Effect of proteolytic inactive versions of the Prc, CtpA and RseP proteases on CSS activity and FoxR RIP. *P. aeruginosa* PAO1 wild-type strain and its isogenic Δprc, ΔctpA and ΔrseP mutants bearing a pBBR1MCS-5 derivative plasmid expressing the indicated *P. aeruginosa* protein and protein variant (-, empty plasmid) (Table S1) were grown under iron-restricted conditions without (-) or with (+) 1 μM ferrioxamine. In **(A)** strains also bear the *P. aeruginosa foxA::lacZ* fusion gene. β-galactosidase activity was determined as described in Materials and Methods and data are means ± SD from three biological replicates (N=3). P-values were calculated by two-tailed *t*-test by comparing the value obtained in the mutant bearing the different protein variants with that obtained in the PAO1 wild-type strain in the same growth condition. Comparisons between other strains are indicated by brackets. In **(B)** strains express an N-terminally HA-tagged *P. aeruginosa* FoxR protein (HA-PaFoxR). Proteins were immunoblotted for HA using a monoclonal antibody. Position of the protein fragments and the molecular size marker (in kDa) is indicated. Presence of the HA-tag adds ~1 kDa to the molar mass of the protein fragments. Blots are representative of at least three biological replicates (N=3).

Next, we assayed the effect of the proteolytically inactive versions of the three proteases on FoxR cleavage by Western-blot analyses. *P. aeruginosa* ΔctpA, Δprc and ΔrseP mutants producing an N-terminally HA-tagged FoxR protein and complemented with either the wild-type or the proteolytically inactive version of the protease were grown in absence and presence of ferrioxamine B (Fig. 4B). The FoxR^N^ domain (~21 kDa) was detected in similar amounts in all strains and conditions analysed (Fig. 4B). Ferrioxamine induction produced the FoxR^ASD^ (~12 kDa) (Fig. 4B, PAO1 strain) as a result of the RIP of the FoxR^N^ domain [11]. As observed before (Fig. 3A), this band was considerably more intense in the ΔctpA mutant, a phenotype that could be complemented with the wild-type CtpA protein but not with the inactive version CtpA-S302A (Fig. 4B, left panel). These results are consistent with the Fox CSS activity observed in these strains (Fig. 4A) relating higher amounts of FoxR^ASD^ with increased CSS activity. Analysis of the FoxR cleavage in the Δprc mutant showed that the band corresponding to the FoxR^ASD^ fragment was slightly less intense than in the PAO1 wild-type strain (Fig. 4B, middle panel), as also observed before (Fig. 3A). However, this phenotype could not be restored upon complementation with the wild-type Prc protein and the amount of FoxR^ASD^ fragment in this strain was also lower than in the PAO1 strain (Fig. 4B, middle panel). In accordance, activity of the Fox CSS system was only partially restored in the complemented Δprc mutant (Fig. 4A). Interestingly, complementation with the proteolytically inactive version of Prc (PaPrc-S490L) resulted in the appearance of a new FoxR fragment of approximately ~19 kDa (FoxR^N1^) (Fig. 4B, middle panel). This band was also detected in the ΔrseP mutant complemented with the PaRseP-H21A variant but not with the wild-type RseP protein (Fig. 4B, right panel). This protein fragment is larger than the FoxR product that accumulates in the ΔrseP mutant (FoxR^N2^, ~15 kDa), which is the substrate of the RseP protease [11]. Presence of FoxR^N1^ suggests that next the site-1 and site-2 cleavages (Fig. 1) another proteolytic event takes place in response to ferrioxamine. Accumulation of FoxR^N1^ only in the Δprc and ΔrseP mutants complemented with the proteolytically inactive version of the protease but not in the other strains suggests that Prc-S390L and RseP-H21A, by being inactive, impede the function of the protease that generates FoxR^N1^ likely by interacting with this protease or its substrate.

### *Lack of RseP, Prc and CtpA proteases affects* P. aeruginosa *virulence*

The CtpA protease has been shown to be required for *P. aeruginosa* virulence in a mouse model of acute infection [15], but the role of RseP and Prc in virulence was not assayed yet. To analyse the pathogenicity of *P. aeruginosa* we used zebrafish (*Danio rerio*) embryos, which are lethally infected by this pathogen when the amount of bacterial cells injected exceeds the phagocytic capacity of the embryo [21–24]. PAO1 wild-type strain and protease mutants were injected into the bloodstream of one-day old embryos to generate a systemic infection and embryo survival was monitored during five days (Fig. 5A). The ΔctpA mutant showed reduced virulence in zebrafish embryos (P<0.0001) (Fig. 5A), in accordance with the reduced virulence previously observed in mice [15]. The survival of the embryos injected with the ΔrseP mutant was similar to that of the control group injected only with phosphate-free physiological salt showing that this mutant was completely attenuated for virulence (P< 0.0001) (Fig. 5A). In contrast, deletion of *prc* resulted in considerably more virulent strain than the PAO1 wild-type strain (P<0.0001) (Fig. 5A). We also used the A549 human respiratory epithelial cell line as host since *P. aeruginosa* often colonizes the human respiratory tract. The cytotoxicity of *P. aeruginosa* towards the eukaryotic cells was determined by measuring A549 cell viability after co-incubation with the bacteria. The ΔrseP mutant was less efficient in damaging the A549 cells than the PAO1 wild-type strain, while the cytotoxicity of the Δprc mutant was similar to that of the wild-type strain (Fig. 5B). In accordance, time-lapse imaging showed that the A549 cells detached after co-incubation with the wild-type strain but not with the ΔrseP mutant (Video 1). In fact, the percentage of detached cells was considerably lower following infection with the mutant (P<0.001) (Fig. S3). Detached is the first indication of cell death. All together, these results indicate that lacks of RseP produces a considerably less virulent *P. aeruginosa* strain.

**Fig. 5.**
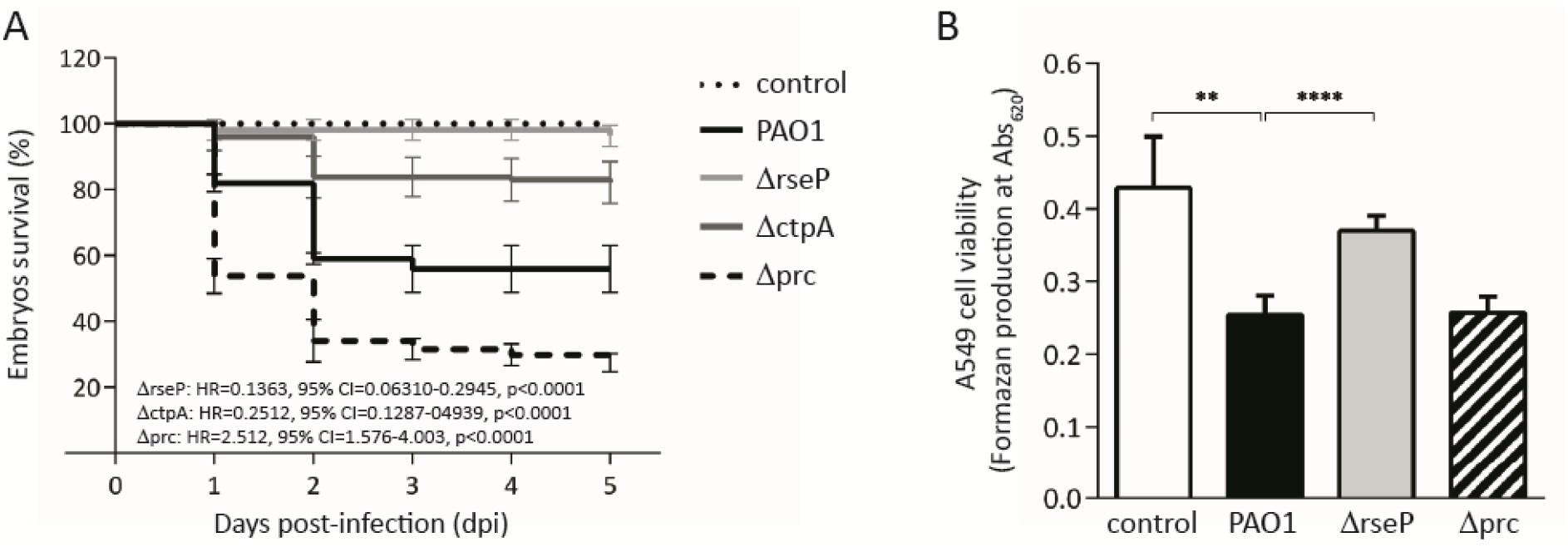
*P. aeruginosa* infections in zebrafish embryos and in the A549 cell line. **(A)** Klapan-Meier zebrafish embryo survival curves upon infection with *P. aeruginosa*. One day-old embryos were injected with ~700-1000 CFU of the *P. aeruginosa* PAO1 wild-type strain or with the indicated isogenic mutant. Uninfected control (non-injected) is shown. Data are means ± SD of five biologically independent replicates (N=5) with 20 embryos/group in each replicate. The hazard ratio (HR), the 95% confidence interval ratio (95% CI), and the p-values (log-rank Mantel-Cox test) of each curve with respect to the reference PAO1 curve (HR=1) are indicated. **(B)** A549 cell viability. The *P. aeruginosa* PAO1 wild-type strain and the indicated isogenic mutant were co-incubated with the eukaryotic cells. Formazan production upon addition of the MTT tretazolium salt was determined spectrophotometrically at 620 nm. Uninfected cells (white bar) were used as control. Data are means ± SD from five biological replicates (N=5). P-values were calculated by two-tailed *t*-test and brackets indicate the comparison to which the P-value applies.

### *Prc modulates the production of* P. aeruginosa *outer membrane vesicles (OMVs)*

An unexpected result from the virulence assays was the hyper-virulent phenotype of the Δprc mutant in zebrafish embryos (Fig. 5). In *E. coli*, the Prc protease (also known as Tsp) modulates the activity of the murein DD-endopeptidase MepS, a hydrolase that cleaves peptidoglycan cross-links to insert new material [25]. The absence of Prc increases MepS levels and uncontrolled MepS activity increases the formation of outer membrane vesicles (OMVs) [26]. Because secretion of OMVs is used by *P. aeruginosa* to deliver multiple virulence factors directly into the host cell cytoplasm [27, 28], we hypothesized that the hyper-virulent phenotype of the Δprc mutant could be related with increased OMV production. SDS-PAGE and transmission electron microscopy (TEM) analyses showed that OMVs production was indeed considerably higher in the Δprc mutant than in the PAO1 wild-type strain, a phenotype that could be complemented by providing the *prc* gene to the mutant in *trans* (Fig. 6A and 6B). This indicates that the absence of Prc raises OMV secretion, which would indeed increase *P. aeruginosa* virulence.

**Fig. 6.**
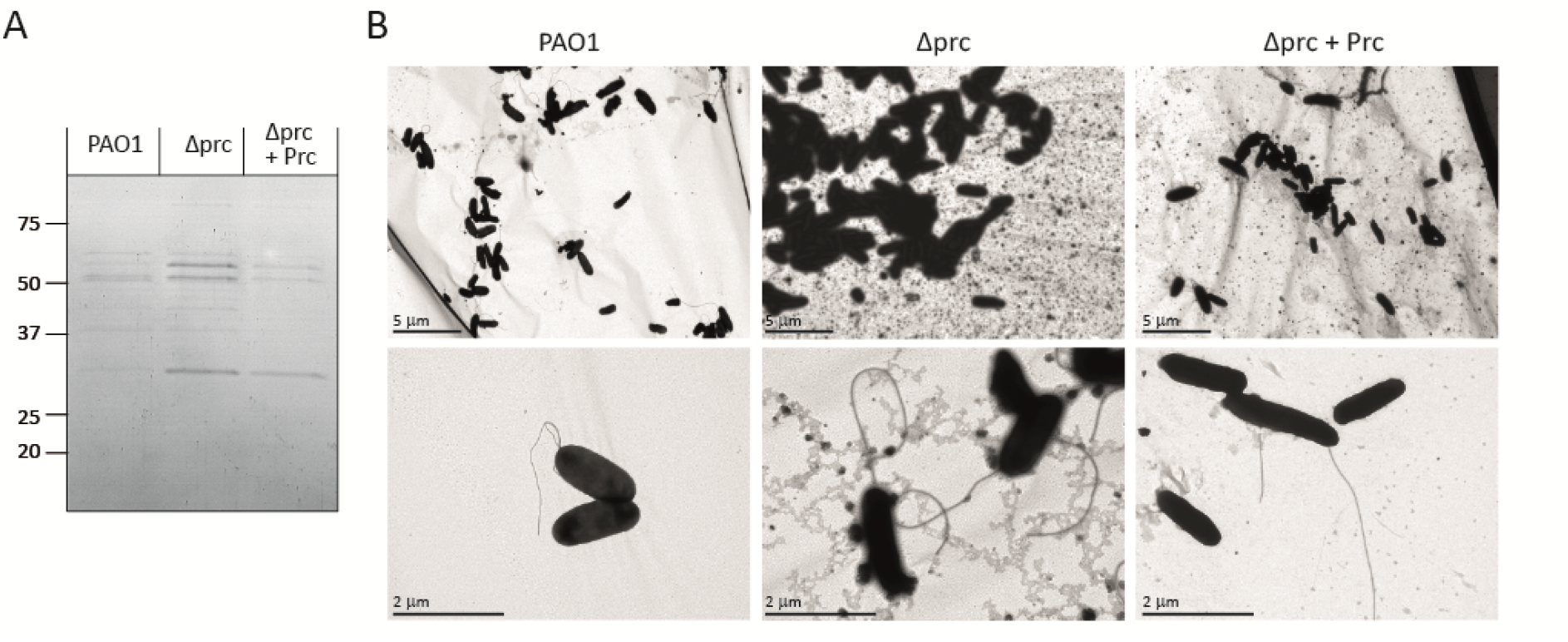
*P. aeruginosa* outer membrane vesicles (OMVs) production and role of the Prc protease. *P. aeruginosa* PAO1 wild-type strain, the isogenic Δprc mutant, and the Δprc mutant bearing the pBBR/PAPrc plasmid (Table S1) (Δprc + Prc) were grown to exponential phase. **(A)** Outer membrane proteins were isolated and visualized by Coomassie staining. Molecular size marker (in kDa) is indicated. SDS-PAGE is representative of three biological replicates (N=3). **(B)** Strains were negatively stained with phosphotungstic acid. TEM images were taken in several fields at 6.000 × and 25.000 ×. Images are representative of three biological replicates (N=3).

## Discussion

In recent years, it has become clear that regulated proteolysis is an important post-translational modification controlling the activity of several signal transduction pathways in bacteria [29]. Signal response proteins are often produced in an inactive state that prevents their interaction with the RNAP and/or the DNA in absence of the inducing signal. Because proteolysis is a fast and irreversible process, it allows for an immediate and longer response than other post-translational modification mechanisms. This may be beneficial in the control of processes like iron homeostasis, stress responses, development or virulence [29]. Moreover, regulated intramembrane proteolysis (RIP) has become increasingly recognized as one of the solution to transmit extracytosolic signals to the cytosol through the cytoplasmic membrane [8]. By this mechanism, proteins are cleaved within the plane of the membrane liberating a cytosolic domain or protein able to modify gene transcription and thus elicit a response. The involvement of RIP in the control of *P. aeruginosa* σ^ECF^ factors activity was first described for σ^AlgT^, which promotes alginate production and the clinical relevant mucoid phenotype of *P. aeruginosa* [30]. Two proteases are required for σ^AlgT^ activation through RIP of is cognate membrane-embedded anti-σ factor MucA: the site-1 protease DegS (also known as AlgW) that also functions as the sensor protein of the signalling pathway and the site-2 protease RseP (also named MucP). In contrast to σ^AlgT^/MucA, CSS σ^ECF^/anti-σ factor pairs usually associate with an outer membrane receptor, which is the component of the signalling pathway that senses the inducing signal [1, 2] (Fig. 1). It is thus not surprising that DegS is not involved in activation of CSS σ^ECF^ factors [10]. RseP/MucP is also the site-2 protease of CSS anti-σ factors [9–12]. In accordance, inactivation of the proteolytic function of RseP completely blocks the activation of *Pseudomonas* CSS pathways (Fig. 4A and S2). For RseP being able to cleave the CSS anti-σ factor, the anti-σ factor needs to first be cleaved by a site-1 protease that generates a smaller form of this protein suitable to enter the catalytic site of RseP [7, 10, 11]. The CTP serine protease Prc was the first identified protease involved in the site-1 cleavage of CSS anti-σ factors, specifically the hybrid σ^ECF^/anti-σ factor protein IutY. In agreement, lack of Prc completely blocks activation of the Iut signalling pathway [9, 10]. However, the activity of the Fox and Fiu CSS systems, in which the σ^ECF^ and anti-σ factor functions reside in two different proteins, is not completely null in a *prc* mutant (Fig. 2 and [10, 11]). Interestingly, a proteolytically inactive version of Prc blocks the activity of CSS systems to a larger extend than the *prc* mutation in both *P. aeruginosa* and *P. putida* (Fig. 4A and S2, Prc-S490L and Prc-S485A proteins, respectively). This confirms that the proteolytic activity of Prc is required to activate the CSS pathway. Moreover, the dominant negative effect exerted by the proteolytically inactive protein is likely the result of tight binding but not cleavage of the protease to its substrate, which avoids other proteolytic events to happen. Prc seems to be active under non-CSS inducing conditions [9], which suggests that Prc function in absence of the signal is blocked by another element. This role could be performed by the CtpA protease, because lack of this protein increases CSS activation in a Prc-dependent manner (Fig. 2). Both Prc and CtpA belong to the S41 family of CTP serine proteases [31], which contain a PDZ domain located upstream the catalytic site required for substrate recognition and activity regulation [18]. *P. aeruginosa* Prc and CtpA are not orthologues; they have only 34% identity, differ in size (Prc is ~76 kDa and CtpA ~44 kDa) and belong to different CTP groups (Prc to the CTP-1 and CtpA to the CTP-3 group) [32]. Western-blot analyses of the *P. aeruginosa* FoxR anti-σ factor, which undergoes spontaneous cleavage immediately upon production that separates the protein in a FoxR^N^ and a FoxR^C^ domain (Fig. 1A), indicate that both Prc and CtpA modulate the levels of the FoxR^C^ domain. Lack of CtpA reduces while that of Prc increases the stability of this domain (Fig. 3B). This suggests that Prc degrades FoxR^C^ while CtpA prevents this degradation to occur. CtpA could perform this function by interacting directly with FoxR^C^ protecting it from degradation or by inhibiting Prc function. We observed reduced instead of increased Prc protein level in the *ctpA* mutant (Fig. 3C), which indicates that CtpA does not degrade Prc. Still, CtpA could act on another element required for Prc function. Interestingly, some CTP proteases partner with a lipoprotein that enhance the proteolysis process. For example, *P. aeruginosa* CtpA partners with the outer membrane lipoprotein LbcA [19, 33] while the *E. coli* Prc protease associates with the NlpI lipoprotein [34, 35]. Because several substrates of the *P. aeruginosa* CtpA protease are outer membrane lipoproteins [19], CtpA could inhibit the function of Prc by degrading its partnering lipoprotein. The lipoprotein that associates with *P. aeruginosa* Prc has not been identified yet and further research is needed to test this hypothesis.

Deleting *ctpA* and *prc* does not prevent the RIP of the *P. aeruginosa* FoxR^N^ domain upon signal recognition (Fig. 3A). Therefore, none of these proteases seems to be the site-1 protease that initiates the RIP pathway of the FoxR^N^ domain in response to ferrioxamine (Fig. 1B) (Bastiaansen et al., 2014; Bastiaansen et al., 2015b). However, absence of CtpA increases the amount of the FoxR RIP product, the FoxR^ASD^ (Fig. 3A). This effect is likely due to the lack of the FoxR^C^ domain in the *ctpA* mutant (Fig. 3B) because it has been shown that FoxR^C^ protects FoxR^N^ from the RIP cascade [11]. In fact, when the FoxR^C^ domain is absent, the activity of the Fox CSS pathway increases [11], as also observed in the *ctpA* mutant (Fig. 2A). Together, these results indicate that CtpA fine-tunes the activity of the *P. aeruginosa* Fox CSS pathway by preventing degradation and thus maintaining the correct levels of the FoxR^C^ anti-σ factor domain, which in turn prevents the RIP of the FoxR^N^ domain.

σ^ECF^-mediated signalling controls important virulence functions in *P. aeruginosa*, including iron acquisition during infection, the response to the oxidative and cell envelope stress produced by components of the immune system of the host, synthesis of the exopolysaccharide alginate responsible of the virulent mucoid phenotype of *P. aeruginosa*, biofilm formation, and production of several virulent determinants (i.e. toxins, exoproteases, secretion systems and secreted proteins) [2, 14, 22, 36, 37]. Therefore, we hypothesized that the proteases required for σ^ECF^ factor activation could be involved in *P. aeruginosa* virulence. Importantly, our results show that a mutant in the RseP protease, which is required for the activation of *P. aeruginosa* σ^ECF^ factors associated with transmembrane anti-σ factors [1, 2, 7, 37], is completely attenuated for virulence and shows significantly reduced cytotoxicity toward host cells (Fig. 5 and Video 1). Although we cannot rule out the possibility that this effect may be due to the role of RseP in processing other transmembrane proteins, we believe that the inability of the *rseP* mutant to carry out σ^ECF^-signalling also contributes to this phenotype. Furthermore, mutation of the CtpA protease reduced *P. aeruginosa* virulence in zebrafish embryos, as other authors observed in mice [15]. This phenotype is likely due to the impaired functioning of the type 3 secretion system (T3SS), which *P. aeruginosa* uses to inject toxic effectors into eukaryotic cells [15]. Surprisingly, mutation of the Prc protease produces a *P. aeruginosa* hyper-virulent strain in zebrafish embryos. In contrast, the *prc* mutation decreases the ability of a pathogenic *E. coli* strain to cause bacteraemia and increases its sensitivity to complement-mediated serum killing [38]. The higher virulence of the *P. aeruginosa prc* mutant could be related to the increased production of outer membrane vesicles (OMVs) observed in this mutant (Fig. 6). *P. aeruginosa* OMVs are known to transport virulence factors to host cells and to promote inflammatory responses [27, 39, 40]. Further analyses will confirm this relation.

In summary, we report in this work the characterization of three proteases that directly or indirectly participate in the activation of *P. aeruginosa* σ^ECF^ factors, shedding more light on the complex proteolytic pathway that controls this process. Importantly, blocking signalling mechanisms required for crucial processes such as iron acquisition is an interesting strategy for drug development that would prevent pathogens from colonizing the host [41]. Proteases are druggable proteins and therefore proteases that modulate the activity of signalling systems required for pathogen’s survival represent excellent drug targets. In fact, an inhibitor of the site-2 RseP protease was already shown to considerably decrease *E. coli* survival (Konovalova et al., 2018). Identification and characterization of new regulatory proteases involved in bacterial signal transfer processes thus holds promise for the development of novel antibacterials.

## Methods

### Bacterial strains and growth conditions

Strains used in this study are listed in Table S1. Construction of the ΔctpA mutants was performed by allelic exchange using a derived of the suicide vector pKNG101, the pKΔctpA plasmid, as described before [10]. Southern blot analyses to confirm the chromosomal gene deletion were performed as described [42]. Bacteria were routinely grown in liquid LB (Sambrook J 1989) on a rotatory shaker at 37 °C and 200 rpm. For low iron conditions, cells were cultured in CAS medium [16] containing 400 μM (for *P. aeruginosa*) or 200 μM (for *P. putida*) of 2,2’-bipyridyl. For induction experiments the low iron media was supplemented with 1 μM of ferrioxamine B (Sigma-Aldrich) or 40 μM of iron-free ferrichrome (Santa Cruz Biotechnology). When required, 1 mM isopropyl β-D-1-thiogalactopyranoside (IPTG) was added to the medium to induce full expression from the pMMB67EH P*tac* promoter. Antibiotics were used at the following final concentrations (μg/ml): ampicillin (Ap), 100; gentamycin (Gm), 10; kanamycin (Km), 100; nalidixic acid (Nal), 25; piperacillin (Pip), 25; streptomycin (Sm), 100; tetracycline (Tc), 20.

### Plasmid construction and molecular biology

Plasmids used are described in Table S1 and primers listed in Table S2. PCR amplifications were performed using Phusion^®^ Hot Start High-Fidelity DNA Polymerase (Finnzymes) or Expand High Fidelity DNA polymerase (Roche). Chromosomal DNA from PAO1 or KT2440 was normally used as DNA template in PCR reactions except for the amplification of the *P. aeruginosa* proteolytically inactive versions of the proteases in which the pBBR1MCS-5 derivative plasmid bearing the wild-type version of the protease gene was used as template (pBBR/PaCtpA, pBBR/PaPrc and pBBR/PaRseP, respectively). All constructs were confirmed by DNA sequencing and transferred to *P. aeruginosa* or *P. putida* by electroporation [43].

### β-galactosidase activity assay

β-galactosidase activities in soluble cell extracts were determined using *o*-nitrophenyl-b-D-galactopyranoside (ONPG) (Sigma-Aldrich) as described before [16]. Each condition was tested in duplicate in at least three biologically independent experiments and the data given are the average of the three biologically independent experiments with error bars representing standard deviation (SD). Activity is expressed in Miller units.

### SDS-PAGE and Western-blot

*P. aeruginosa* was grown in iron-limited medium containing 1 mM IPTG (when necessary) and without or with 1 μM ferrioxamine. Cells were pelleted by centrifugation and heated for 10 min at 95 °C following solubilisation in SDS-PAGE sample buffer [44]. Sample normalization was done according to the OD_660_ of the bacterial culture. Proteins were separated by SDS-PAGE containing 8, 12 or 15% (w/v) acrylamide and electrotransferred to nitrocellulose membranes (Millipore). Ponceau S (Serva) staining was performed as a loading control. Immunodetection was realized using monoclonal antibody directed against the influenza hemagglutinin epitope (HA.11, Covance, Princeton, NJ) or polyclonal antibodies directed against the *P. aeruginosa* OprF [45] or FoxA proteins. The FoxA antibody was generated at Abyntek using the peptide SDTQFDHVKEERYAC as antigen. The second antibody, either the horseradish peroxidase-conjugated rabbit anti-mouse (DAKO) or the horseradish peroxidase-conjugated goat anti-rabbit IgG (Sigma-Aldrich), was detected using the SuperSignal^®^ West Femto Chemiluminescent Substrate (Thermo Scientific). Blots were scanned and analysed using the Quantity One version 4.6.7 (Bio-Rad).

### Zebrafish maintenance, embryo care and infection procedure

Transparent adult *casper* mutant zebrafish (*mitfa ^w2/w2^;roy ^a9/a9^*) [46, 47] were conserved at 26 °C in aerated 5 L tanks with a 10/14 h dark/light cycle. Zebrafish embryos were collected during the first hour post-fertilization (hpf) and kept at 28 °C in E3 medium (5.0 mM NaCl, 0.17 mM KCl, 0.33 mM CaCl·2H_2_O, 0.33 mM MgCl_2_·7H_2_O) supplemented with 0.3 mg/L methylene blue. Prior to infection, 1day post-fertilization (dpf) embryos were mechanically dechorionated and anaesthetized in 0.02 % (*w/v*) buffered 3-aminobenzoic acid methyl ester (pH 7.0) (Tricaine, Sigma-Aldrich). Zebrafish embryos were individually infected by microinjection with 1 nl of *P. aeruginosa* in the caudal vein (systemic infection) as described elsewhere [23]. All procedures involving zebrafish embryos were according to local animal welfare regulations.

### Virulence assay in infected zebrafish embryos

Zebrafish embryos were injected in the caudal vein with ~1000 CFU of exponentially grown *P. aeruginosa* cells previously suspended in phosphate-free physiological salt containing 0.5 % (w/v) of phenol red. After infection, embryos were kept in 12-well plates containing 60 μg/mL of Sea salts (Sigma-Aldrich) at 32 °C with 20 individually injected embryos in each group per well. Embryo survival was determined by monitoring live and dead embryos at fixed time points during five days. Five biologically independent experiments were performed and the data given are the average. P-values were calculated by log-rank (Mantel-Cox) test.

### Cytotoxicity assay in A549 human lung epithelial cells

*P. aeruginosa* cytotoxicity on A549 cells was assayed using a colorimetric assay that detects the number of metabolically active eukaryotic cells able to cleave the MTT tetrazolium salt (Sigma-Aldrich) to the insoluble formazan dye. The A549 cell line (ATCC® CCL-185™) was maintained in DMEM medium supplemented with 10 % (v/v) fetal bovine serum (FBS) (Gibco) in a 5 % CO_2_ incubator at 37 °C. One-day prior infection, the A549 cells were placed in 96-well plates at a concentration of 4 × 10^4^ cells/well and cultured in phosphate-free DMEM medium (Gibco) with 5 % (v/v) FBS. In this condition, cell mitosis does almost not occur. Late exponentially grown *P. aeruginosa* strains were then inoculated at a multiplicity of infection (MOI) of 20. At 3 hpi, 30 μl of a 5 mg/ml MTT solution in PBS was added to the wells and the plates were incubated for 2 h. The culture medium was then removed and 100 μl of dimethyl sulfoxide (DMSO) was added to solubilize the formazan. Production of formazan, which directly correlates to the number of viable cells, was quantified using a scanning multi-well spectrophotometer (Infinite® 200 PRO Tecan) at 620 nm.

### Time-lapse imaging assay

Time-lapse microscopy was performed on a Nikon Eclipse Ti-E microscope (Nikon), equipped with a PlanFluor 20–40 × 0.6NA objective (Nikon) and a CO_2_ incubator. A549 cells were seeded in coated 4-well μ-slides (Ibidi, Martinsried, Germany) in the same conditions as in the cytotoxic assays. Late exponentially grown *P. aeruginosa* strains were then inoculated at a multiplicity of infection (MOI) of 20. Images were collected from 0 to 240 min post-infection every 2 min with an ORCA- R2 CCD camera (Hamamatsu) powered by Nis Elements 3.2 software. Videos where edited with ImageJ.

### Outer membrane vesicles (OMVs) isolation and detection

*P. aeruginosa* OMVs were isolated following the protocol described before [48] with some modifications. Briefly, overnight LB cultures of *P. aeruginosa* were diluted at 0.05 OD_600_. At OD_600_ of ~1, the cultures were collected and centrifuged at 6.000 × g, 4 °C. Approximately 10 ml of the supernatants were filtered through a 0.45 μm pore size filter (Millipore) to remove non-pelleted cells and the OMVs were pelleted by ultracentrifugation at 150.000 × g, 180 min, 4 °C. After carefully removal of the supernatant, the pellet was resuspended in PBS containing 15 % (v/v) glycerol in a volume proportional to the final OD_600_ of the culture and the amount of supernatant ultracentrifuged (e.g. 10 ml of supernatant of a 1.2 OD_600_ culture was resuspended in 120 μl of PBS-glycerol). 5 μl of the suspension was plated to verify that it was free of bacteria. OMV proteins (40 μl) were separated by SDS-PAGE containing 15 % (w/v) acrylamide and visualized by Coomassie staining.

### Transmission electron microscopy (TEM)

TEM samples were prepared following the method described before [49]. Overnight cultures of *P. aeruginosa* were centrifuged at 0.6 × g and the pellets were washed four times in PBS. Bacteria cells were placed in 100 mesh copper grids containing the support carbon-coated Formvar film (Electron Microscopy Sciences) and air dried. Cells were negatively stained, with 1 % (w/v) phosphotungstic acid in distilled water, for 20 secs and examined with a JEOL JEM-1011 transmission electron microscope equipped with an ORIUS SC 1000 CCD camera (GATAN).

### Computer-assisted analysis

Images processing and band intensity measurements were performed using the ImageJ software. Statistical analyses are based on *t*-test in which two conditions are compared independently. P-values from raw data (i.e. miller units from β-galactosidase assays) were calculated by independent two-tailed *t*-test, from ratio data to the control (i.e. FoxA production in Fig. 2C) by one-sample *t*-test, and Kaplan-Meyer survival curves by Log-Rank (Mantel-Cox) using GRAPHPAD PRISM version 5.01 for Windows and are represented in the graphs by ns, not significant; *, P<0.05; **, P<0.01; ***, P<0.001; and ****, P<0.0001.

## Acknowledgments

We thank A. Ocampo and F. Madrazo (Hospital Universitario Marqués de Valdecilla) for assistance with A549 cytotoxicity assays and time-lapse imaging, and K. K. Jim for assistance with the zebrafish embryo infections.

## Author contributions

JOA, WB and MAL designed the study. JOA, KB, ASJ, SW, CC and AGP performed experiments. JOA, WB and MAL contributed to the interpretation of the results. JOA and MAL wrote the manuscript with input from SW and WB.

## Additional Information

The authors declare that they have no competing interests.

## References

1. Otero-Asman JR, Wettstadt S, Bernal P, Llamas MA. Diversity of extracytoplasmic function sigma (σ^ECF^) factor-dependent signaling in *Pseudomonas*. Mol Microbiol. 2019;112(2):356–73. doi: 10.1111/mmi.14331. PubMed PMID: 31206859.

2. Llamas MA, Imperi F, Visca P, Lamont IL. Cell-surface signaling in *Pseudomonas*: stress responses, iron transport, and pathogenicity. FEMS Microbiol Rev. 2014;38(4):569–97. doi: 10.1111/1574-6976.12078. PubMed PMID: 24923658.

3. Noinaj N, Guillier M, Barnard TJ, Buchanan SK. TonB-dependent transporters: regulation, structure, and function. Annu Rev Microbiol. 2010;64:43–60. doi: 10.1146/annurev.micro.112408.134247. PubMed PMID: 20420522; PubMed Central PMCID: PMCPMC3108441.

4. Enz S, Brand H, Orellana C, Mahren S, Braun V. Sites of interaction between the FecA and FecR signal transduction proteins of ferric citrate transport in *Escherichia coli* K-12. J Bacteriol. 2003;185(13):3745–52. PubMed PMID: 160.

5. Spiers AJ, Buckling A, Rainey PB. The causes of *Pseudomonas* diversity. Microbiology. 2000;146:2345–50. Epub 2000/10/06. PubMed PMID: 11021911.

6. Silby MW, Winstanley C, Godfrey SA, Levy SB, Jackson RW. *Pseudomonas* genomes: diverse and adaptable. FEMS Microbiol Rev. 2011;35(4):652–80. Epub 2011/03/03. doi: 10.1111/j.1574-6976.2011.00269.x. PubMed PMID: 21361996.

7. Otero-Asman JR, Garcia-Garcia AI, Civantos C, Quesada JM, Llamas MA. *Pseudomonas aeruginosa* possesses three distinct systems for sensing and using the host molecule haem. Environ Microbiol. 2019;21(12):4629–47. doi: 10.1111/1462-2920.14773. PubMed PMID: 31390127.

8. Brown MS, Ye J, Rawson RB, Goldstein JL. Regulated intramembrane proteolysis: a control mechanism conserved from bacteria to humans. Cell. 2000;100(4):391–8. PubMed PMID: 10693756.

9. Bastiaansen KC, Civantos C, Bitter W, Llamas MA. New insights into the regulation of cell-surface signaling activity acquired from a mutagenesis screen of the *Pseudomonas putida* IutY sigma/anti-sigma factor. Front Microbiol. 2017;8-747(747):1–15. doi: 10.3389/fmicb.2017.00747. PubMed PMID: 28512454; PubMed Central PMCID: PMCPMC5411451.

10. Bastiaansen KC, Ibañez A, Ramos JL, Bitter W, Llamas MA. The Prc and RseP proteases control bacterial cell-surface signalling activity. Environ Microbiol. 2014;16(8):2433–43. Epub 2014/01/01. doi: 10.1111/1462-2920.12371. PubMed PMID: 24373018.

11. Bastiaansen KC, Otero-Asman JR, Luirink J, Bitter W, Llamas MA. Processing of cell-surface signalling anti-sigma factors prior to signal recognition is a conserved autoproteolytic mechanism that produces two functional domains. Environ Microbiol. 2015;17(9):3263–77. Epub 2015/01/13. doi: 10.1111/1462-2920.12776. PubMed PMID: 25581349.

12. Draper RC, Martin LW, Beare PA, Lamont IL. Differential proteolysis of sigma regulators controls cell-surface signalling in *Pseudomonas aeruginosa*. Mol Microbiol. 2011;82(6):1444–53. doi: 10.1111/j.1365-2958.2011.07901.x. PubMed PMID: 22040024.

13. Hizukuri Y, Oda T, Tabata S, Tamura-Kawakami K, Oi R, Sato M, et al. A structure-based model of substrate discrimination by a noncanonical PDZ tandem in the intramembrane-cleaving protease RseP. Structure. 2014;22(2):326–36. doi: 10.1016/j.str.2013.12.003. PubMed PMID: 24389025.

14. McGuffie BA, Vallet-Gely I, Dove SL. σ factor and anti-σ factor that control swarming motility and biofilm formation in *Pseudomonas aeruginosa*. J Bacteriol. 2015;198(5):755–65. Epub 2015/12/02. doi: 10.1128/JB.00784-15. PubMed PMID: 26620262; PubMed Central PMCID: PMCPMC4810613.

15. Seo J, Darwin AJ. The *Pseudomonas aeruginosa* periplasmic protease CtpA can affect systems that impact its ability to mount both acute and chronic infections. Infect Immun. 2013;81(12):4561–70. doi: 10.1128/IAI.01035-13. PubMed PMID: 24082078; PubMed Central PMCID: PMCPMC3837984.

16. Llamas MA, Sparrius M, Kloet R, Jimenez CR, Vandenbroucke-Grauls C, Bitter W. The heterologous siderophores ferrioxamine B and ferrichrome activate signaling pathways in *Pseudomonas aeruginosa*. J Bacteriol. 2006;188(5):1882–91. PubMed PMID: 2.

17. Bastiaansen KC, van Ulsen P, Wijtmans M, Bitter W, Llamas MA. Self-cleavage of the *Pseudomonas* aeruginosa Cell-surface signaling anti-sigma factor FoxR occurs through an N-O acyl rearrangement. J Biol Chem. 2015;290(19):12237–46. Epub 2015/03/27. doi: 10.1074/jbc.M115.643098. PubMed PMID: 25809487; PubMed Central PMCID: PMC4424355.

18. Chueh CK, Som N, Ke LC, Ho MR, Reddy M, Chang CI. Structural basis for the differential regulatory roles of the PDZ domain in C-Terminal processing proteases. mBio. 2019;10(4). Epub 20190806. doi: 10.1128/mBio.01129-19. PubMed PMID: 31387902; PubMed Central PMCID: PMCPMC6686036.

19. Srivastava D, Seo J, Rimal B, Kim SJ, Zhen S, Darwin AJ. A proteolytic complex targets multiple cell wall hydrolases in *Pseudomonas aeruginosa*. mBio. 2018;9(4):e00972. doi: doi:10.1128/mBio.00972-18.

20. Koide K, Maegawa S, Ito K, Akiyama Y. Environment of the active site region of RseP, an *Escherichia coli* regulated intramembrane proteolysis protease, assessed by site-directed cysteine alkylation. J Biol Chem. 2007;282(7):4553–60. doi: 10.1074/jbc.M607339200. PubMed PMID: 17179147.

21. Clatworthy AE, Lee JS, Leibman M, Kostun Z, Davidson AJ, Hung DT. *Pseudomonas aeruginosa* infection of zebrafish involves both host and pathogen determinants. Infect Immun. 2009;77(4):1293–303. PubMed PMID: 215.

22. Llamas MA, van der Sar A, Chu BC, Sparrius M, Vogel HJ, Bitter W. A novel extracytoplasmic function (ECF) sigma factor regulates virulence in *Pseudomonas aeruginosa*. PLoS Pathog. 2009;5(9):e1000572. PubMed PMID: 231.

23. Llamas MA, van der Sar AM. Assessing *Pseudomonas* virulence with nonmammalian host: zebrafish. Methods Mol Biol. 2014;1149:709–21. Epub 2014/05/14. doi: 10.1007/978-1-4939-0473-0_55. PubMed PMID: 24818945.

24. Otero-Asman JR, Quesada JM, Jim KK, Ocampo-Sosa A, Civantos C, Bitter W, et al. The extracytoplasmic function sigma factor σ^VreI^ is active during infection and contributes to phosphate starvation-induced virulence of *Pseudomonas aeruginosa*. Sci Rep. 2020;10(1):3139. doi: 10.1038/s41598-020-60197-x. PubMed PMID: 32081993; PubMed Central PMCID: PMCPMC7035377.

25. Singh SK, Parveen S, SaiSree L, Reddy M. Regulated proteolysis of a cross-link-specific peptidoglycan hydrolase contributes to bacterial morphogenesis. Proc Natl Acad Sci U S A. 2015;112(35):10956–61. Epub 2015/08/19. doi: 10.1073/pnas.1507760112. PubMed PMID: 26283368; PubMed Central PMCID: PMCPMC4568209.

26. Schwechheimer C, Rodriguez DL, Kuehn MJ. NlpI-mediated modulation of outer membrane vesicle production through peptidoglycan dynamics in *Escherichia coli*. Microbiologyopen. 2015;4(3):375–89. doi: 10.1002/mbo3.244. PubMed PMID: 25755088; PubMed Central PMCID: PMCPMC4475382.

27. Bomberger JM, Maceachran DP, Coutermarsh BA, Ye S, O’Toole GA, Stanton BA. Long-distance delivery of bacterial virulence factors by *Pseudomonas aeruginosa* outer membrane vesicles. PLoS Pathog. 2009;5(4):e1000382. doi: 10.1371/journal.ppat.1000382. PubMed PMID: 19360133; PubMed Central PMCID: PMCPMC2661024.

28. Koeppen K, Hampton TH, Jarek M, Scharfe M, Gerber SA, Mielcarz DW, et al. A novel mechanism of host-pathogen interaction through sRNA in bacterial outer membrane vesicles. PLoS Pathog. 2016;12(6):e1005672. doi: 10.1371/journal.ppat.1005672. PubMed PMID: 27295279; PubMed Central PMCID: PMCPMC4905634.

29. Wettstadt S, Llamas MA. Role of regulated proteolysis in the communication of bacteria with the environment. Front Mol Biosci. 2020;7:586497. doi: 10.3389/fmolb.2020.586497. PubMed PMID: 33195433; PubMed Central PMCID: PMCPMC7593790.

30. Qiu D, Eisinger VM, Rowen DW, Yu HD. Regulated proteolysis controls mucoid conversion in *Pseudomonas aeruginosa*. Proc Natl Acad Sci U S A. 2007;104(19):8107–12. doi: 10.1073/pnas.0702660104. PubMed PMID: 17470813; PubMed Central PMCID: PMCPMC1876579.

31. Rawlings ND, Waller M, Barrett AJ, Bateman A. MEROPS: the database of proteolytic enzymes, their substrates and inhibitors. Nucleic Acids Res. 2014;42(Database issue):D503–9. doi: 10.1093/nar/gkt953. PubMed PMID: 24157837; PubMed Central PMCID: PMCPMC3964991.

32. Hoge R, Laschinski M, Jaeger KE, Wilhelm S, Rosenau F. The subcellular localization of a C-terminal processing protease in *Pseudomonas aeruginosa*. FEMS Microbiol Lett. 2011;316(1):23–30. Epub 2011/01/06. doi: 10.1111/j.1574-6968.2010.02181.x. PubMed PMID: 21204920.

33. Hsu HC, Wang M, Kovach A, Darwin AJ, Li H. *Pseudomonas aeruginosa* C-Terminal processing protease CtpA assembles into a hexameric structure that requires activation by a spiral-shaped lipoprotein-binding partner. mBio. 2022:e0368021. Epub 20220118. doi: 10.1128/mbio.03680-21. PubMed PMID: 35038915; PubMed Central PMCID: PMCPMC8764530.

34. Kim YJ, Choi BJ, Park SH, Lee HB, Son JE, Choi U, et al. Distinct amino acid availability-dependent regulatory mechanisms of MepS and MepM levels in *Escherichia coli*. Front Microbiol. 2021;12:677739. Epub 2021/07/20. doi: 10.3389/fmicb.2021.677739. PubMed PMID: 34276609; PubMed Central PMCID: PMCPMC8278236.

35. Su MY, Som N, Wu CY, Su SC, Kuo YT, Ke LC, et al. Structural basis of adaptor-mediated protein degradation by the tail-specific PDZ-protease Prc. Nat Commun. 2017;8(1):1516. Epub 2017/11/16. doi: 10.1038/s41467-017-01697-9. PubMed PMID: 29138488; PubMed Central PMCID: PMCPMC5686067.

36. Hershberger CD, Ye RW, Parsek MR, Xie ZD, Chakrabarty AM. The *algT(algU)* gene of *Pseudomonas aeruginosa*, a key regulator involved in alginate biosynthesis, encodes an alternative sigma factor (σ^E^). Proc Natl Acad Sci U S A. 1995;92(17):7941–5. PubMed PMID: 7644517.

37. Chevalier S, Bouffartigues E, Bazire A, Tahrioui A, Duchesne R, Tortuel D, et al. Extracytoplasmic function sigma factors in *Pseudomonas aeruginosa*. Biochim Biophys Acta Gene Regul Mech. 2019;1862(7):706–21. doi: 10.1016/j.bbagrm.2018.04.008. PubMed PMID: 29729420.

38. Wang CY, Wang SW, Huang WC, Kim KS, Chang NS, Wang YH, et al. Prc contributes to *Escherichia coli* evasion of classical complement-mediated serum killing. Infect Immun. 2012;80(10):3399–409. doi: 10.1128/IAI.00321-12. PubMed PMID: 22825444; PubMed Central PMCID: PMCPMC3457568.

39. Ellis TN, Kuehn MJ. Virulence and immunomodulatory roles of bacterial outer membrane vesicles. Microbiol Mol Biol Rev. 2010;74(1):81–94. doi: 10.1128/MMBR.00031-09. PubMed PMID: 20197500; PubMed Central PMCID: PMCPMC2832350.

40. Lee J, Kim OY, Gho YS. Proteomic profiling of Gram-negative bacterial outer membrane vesicles: Current perspectives. Proteomics Clin Appl. 2016;10(9-10):897–909. Epub 2016/08/03. doi: 10.1002/prca.201600032. PubMed PMID: 27480505.

41. Sánchez-Jiménez A, Marcos-Torres FJ, Llamas MA. Mechanisms of iron homeostasis in *Pseudomonas aeruginosa* and emerging therapeutics directed to disrupt this vital process. Microb Biotechnol. 2023;doi: 10.1111/1751-7915.14241. Epub 2023/03/02. doi: 10.1111/1751-7915.14241. PubMed PMID: 36857468.

42. Llamas MA, Ramos JL, Rodriguez-Herva JJ. Mutations in each of the *tol* genes of *Pseudomonas putida* reveal that they are critical for maintenance of outer membrane stability. J Bacteriol. 2000;182(17):4764–72. PubMed PMID: 7.

43. Choi KH, Kumar A, Schweizer HP. A 10-min method for preparation of highly electrocompetent *Pseudomonas aeruginosa* cells: application for DNA fragment transfer between chromosomes and plasmid transformation. J Microbiol Methods. 2006;64(3):391–7. Epub 2005/07/01. doi: 10.1016/j.mimet.2005.06.001. PubMed PMID: 15987659.

44. Laemmli UK. Cleavage of structural proteins during the assembly of the head of bacteriophage T4. Nature. 1970;227(5259):680–5. PubMed PMID: 69.

45. Rawling EG, Martin NL, Hancock RE. Epitope mapping of the *Pseudomonas aeruginosa* major outer membrane porin protein OprF. Infect Immun. 1995;63(1):38–42. PubMed PMID: 7806382; PubMed Central PMCID: PMCPMC172954.

46. Renshaw SA, Loynes CA, Trushell DM, Elworthy S, Ingham PW, Whyte MK. A transgenic zebrafish model of neutrophilic inflammation. Blood. 2006;108(13):3976–8. doi: 10.1182/blood-2006-05-024075. PubMed PMID: 16926288.

47. White RM, Sessa A, Burke C, Bowman T, LeBlanc J, Ceol C, et al. Transparent adult zebrafish as a tool for in vivo transplantation analysis. Cell Stem Cell. 2008;2(2):183–9. doi: 10.1016/j.stem.2007.11.002. PubMed PMID: 18371439; PubMed Central PMCID: PMCPMC2292119.

48. Balsalobre C, Silvan JM, Berglund S, Mizunoe Y, Uhlin BE, Wai SN. Release of the type I secreted alpha-haemolysin via outer membrane vesicles from *Escherichia coli*. Mol Microbiol. 2006;59(1):99–112. doi: 10.1111/j.1365-2958.2005.04938.x. PubMed PMID: 16359321.

49. Remuzgo-Martinez S, Lazaro-Diez M, Mayer C, Aranzamendi-Zaldumbide M, Padilla D, Calvo J, et al. Biofilm formation and quorum-sensing-molecule production by clinical isolates of *Serratia liquefaciens*. Appl Environ Microbiol. 2015;81(10):3306–15. doi: 10.1128/AEM.00088-15. PubMed PMID: 25746999; PubMed Central PMCID: PMCPMC4407221.

